# Efficient NGN2-induced neuronal differentiation from human stem cells for DNA repair studies and functional analysis

**DOI:** 10.64898/2026.11.26.769571

**Authors:** Bokun Yin, Phillip R. Woolley, Samantha T. Refvik, Hannah Barnes, Tanya T. Paull

## Abstract

**Background:** Mutations in DNA repair factors are increasingly linked to defects in the central nervous system. Although these factors have been studied extensively in proliferating cancer cells, their roles in post-mitotic neurons in vitro remain less well understood. Murine models have advanced our understanding but often fail to recapitulate human neurodegeneration. Human post-mitotic cells reduce species-specific differences and allow disease-associated mutations to be studied in their native genetic context. However, many differentiation protocols depend on glial co-culture or the use of nucleoside analogs, which cause DNA damage and confound repair measurements.

**New method:** Here we describe an optimized NGN2-mediated protocol that generates high-yield, homogeneous, glia-free neurons from human embryonic stem cells in the absence of DNA damaging agents.

**Results:** Using our modified NGN2 protocol, we observe efficient differentiation and maturation to glutamatergic neurons, verified by immunostaining and transcriptomic analysis, in the absence of non-differentiated cell growth. We also optimize a simple assay combining ReaChR optogenetic stimulation with GCaMP3 calcium imaging to assess neuronal function.

**Comparison with existing methods:** This modified protocol with lentiviral delivery of NGN2 allows for selection of the lentivirus separately from differentiation, facilitating unlimited scaling of the culture and further genetic manipulation, including stocking of selected cells. Doxycycline-induced differentiation and maturation of neurons occurs efficiently, without growth of non-differentiated cells, while also avoiding use of nucleoside analogs.

**Conclusion:** This scalable protocol and rapid functional readout provide an accessible platform for generating and validating human glutamatergic neurons for molecular studies of DNA repair and gene function.

## Introduction

Accumulating pathological evidence links mutations in DNA repair factors to defects in the central nervous system (Ropert et al., 2024). While the functions of these factors have been studied extensively in proliferating cells, particularly cancer cell lines, fewer studies have examined their roles in post-mitotic neurons in vitro. Although murine models have contributed substantially to our understanding of DNA repair protein mechanisms in neurons, they often fail to recapitulate human neurodegeneration, as seen in dominant and recessive forms of cerebellar ataxia (Coarelli et al., 2023; Cui et al., 2024; Rothblum-Oviatt et al., 2016). Using post-mitotic cells with a human genomic background reduces species-specific differences in genome organization, protein interactions, and signaling pathways, thus allowing disease-associated mutations to be studied within their native genetic context (Kim et al., 2021; Nayler and Becker, 2018).

Multiple studies have shown that overexpression of the transcription factor NGN2, a master regulator of neural development, gives rise to glutamatergic neurons (Hulme et al., 2022). While the first developed protocols required co-culture with supportive glial cells to promote neuronal differentiation (Busskamp et al., 2014; Zhang et al., 2013), more recent protocols yield pure neuronal cultures without additional cell types (Shan et al., 2024; Wang et al., 2017). Because our downstream applications involve molecular and omics-level analyses requiring large numbers of cells, we focused on establishing a protocol that allows for expansion of transduced cells before differentiation while still generating a pure neuronal culture free of glial contamination. It is also essential that the differentiation protocol avoids the use of nucleoside analogs such as AraC to inhibit the growth of proliferating cells, since this treatment generates DNA damage and can interfere with DNA repair measurements in post-mitotic neurons. Here we established an optimized NGN2-mediated differentiation protocol that generates high-yield, homogeneous, glia-free neurons from human H9 (WA09) embryonic stem cells. By uncoupling antibiotic selection from NGN2 induction, we improved neuronal viability while maintaining efficient differentiation. The protocol prevents growth of non-neuronal cells yet is comparable in efficiency to neuronal differentiation using nucleoside analogs. We also implemented a simple optogenetic calcium-imaging assay for rapid, single-cell assessment of neuronal activity in the induced neurons. Together, these approaches provide a scalable and accessible platform for generating and functionally validating human neurons for studies of gene function in vitro.

## Results and Discussion

### Modifications of the NGN2 neuronal induction system to facilitate larger-scale differentiation of human neurons in vitro without use of nucleoside analogs

To induce neuronal differentiation, human embryonic stem cells (H9) were transduced with a lentiviral construct encoding an inducible NGN2 transgene as previously described (Zhang et al., 2013). Following transduction, cells were selected for lentivirus integration by incubating with puromycin, which simultaneously initiates differentiation. Neuronal differentiation was performed using a two-stage protocol: initiation, which down-regulates pluripotent cell expression, and maturation, which promotes a neural gene expression program and formation of neuronal networks, adapted from established methods (Fig. 1A,B)(Wu et al., 2021; Zhang et al., 2013). Using a previously published lentiviral TetO-driven NGN2 construct containing a puromycin selection cassette (Addgene 79823)(Ho et al., 2016), doxycycline addition successfully induced NGN2 expression and neurite outgrowth. However, puromycin selection for the NGN2 construct during the differentiation stage resulted in substantial loss of cell viability such that the yield of mature neurons after 14 days was less than 1% relative to the number of stem cells originally plated (Fig. 1C).

**Figure 1.**
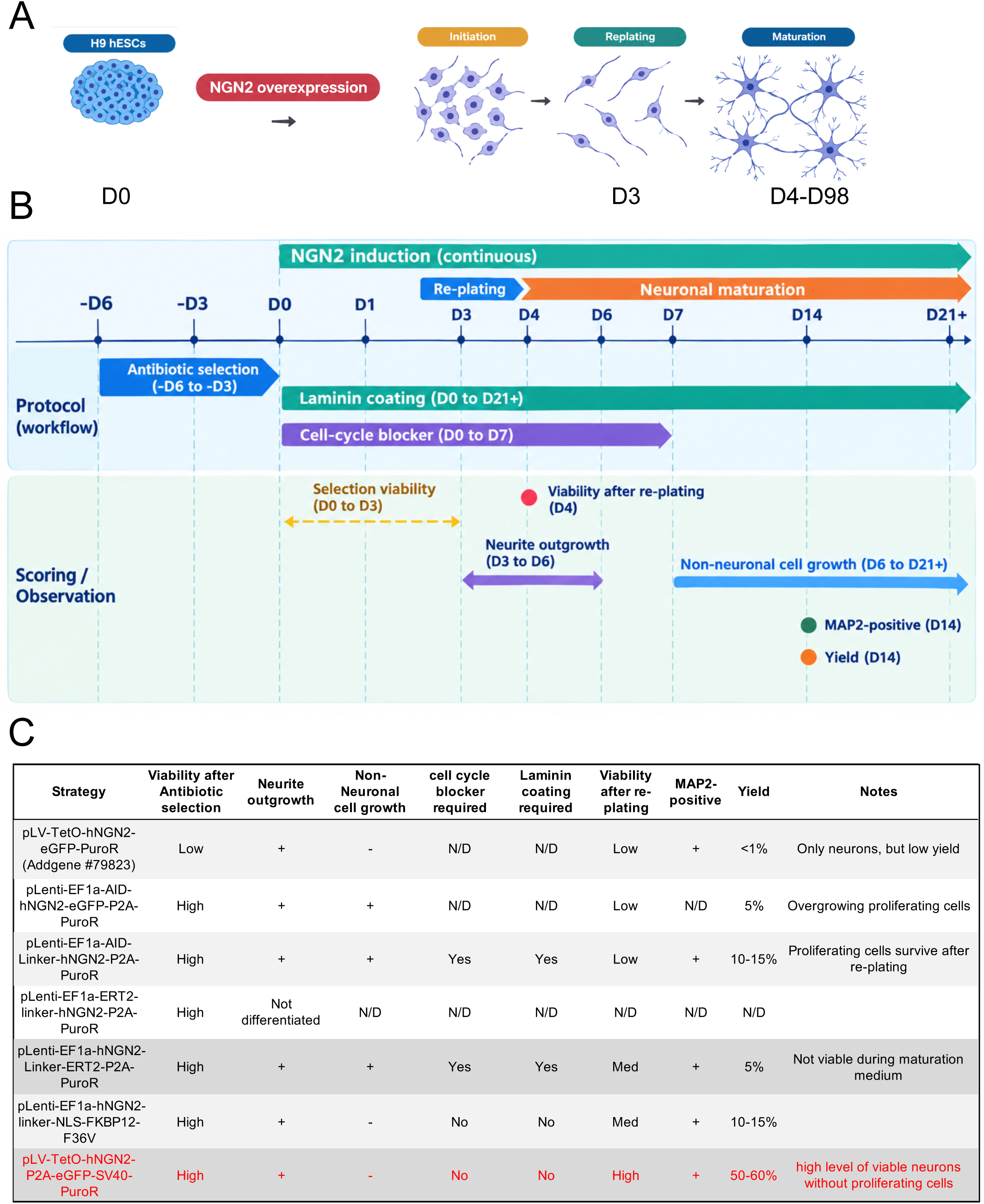
hNGN2-induced differentiation strategies and signs of differentiation. (1A) General schematic of NGN2-induced iNeuron differentiation. After NGN2 overexpression, hESCs initiate neuronal lineage differentiation, followed by replating. Neurons are then further cultured for maturation. (1B) Timeline for scoring neuronal differentiation: antibiotic-selection viability was scored between D-6 and D-3, with the exception of lentiviral TetO-driven NGN2 construct containing a puromycin selection cassette (Addgene 79823) in which cells were selected after induction of differentiation, between D0 and D3. Neurite outgrowth was assessed between D4 and D6, and non-neuronal cell growth after D6. At D14, neuronal yield (%) was scored and the MAP2-positive cells were detected. (1C) Strategies for hNGN2-induced differentiation via lentiviral delivery including the original plasmid (Addgene #79823), alterations containing varying NGN2 fusions, or changes in the regulation of the puromycin resistance cassette. The most successful strategy with puromycin resistance driven by the SV40 promoter is shown in red. Viability was scored after antibiotic selection and after re-plating. For some strategies, laminin coating or cell-cycle blocker 10058-F4 were necessary to support differentiation, as indicated. MAP2 immunostaining was used to identify mature neuronal cells. “Yield” indicates the ratio of scored neurons to the number of H9 cells originally plated. N/D: not determined.

We hypothesized that puromycin selection during active differentiation is detrimental to cell survival. To address this, we tested several lentivirus design strategies in which cells could be selected with puromycin prior to and independent from differentiation induction. These included fusion of NGN2 to auxin-inducible degron (AID)(Nishimura et al., 2009), or regulation of NGN2 via an ERT2-based conditional activation system(Eilers et al., 1989) or an FKBP12(F36V) PROTAC degrader system (Nabet et al., 2018).

The AID system did not yield a pure neuronal culture; proliferating cells persisted and reached confluency during differentiation, even in the presence of the cell-cycle inhibitor 10058-F4 (results summarized in Fig. 1C). The ERT2 system supported neuronal induction, but neurons failed to survive the maturation phase. The FKBP12(F36V) system produced a pure neuronal culture free of proliferating cells; however, overall yield from H9 cells into surviving mature neurons was only 15%.

To improve yield, we uncoupled the puromycin-resistance cassette from the NGN2 cassette, placing puromycin resistance under a constitutive SV40 promoter but retaining tet operator control of NGN2 (pLVTetO-hNGN2-P2A-eGFP SV40-PuroR). Following selection of transduced H9 cells with puromycin, the cells could be expanded in the absence of doxycycline, further manipulated with other viral constructs or treatments, and stocked for later use without initiating the differentiation process. We concluded that this strategy was preferable to others and moved forward with this construct which produced approximately 50-60% overall neuronal yield relative to the number of stem cells originally plated (Fig. 1C).

Our modified strategy in which selection of the NGN2-containing lentivirus with puromycin selection occurs first, followed by doxycycline-induced NGN2-P2A-GFP expression, replating, and maturation in the absence of nucleoside analogs or cell cycle blockers is summarized in Fig. 2. The protocol includes the same steps as the original protocols, including induction of differentiation followed by maturation and replating of cells on Poly-L-Ornithine-coated plates in presence of Brain Phys neuronal media (Bardy et al., 2015) supplemented with B-27, NT-3, BDNF, and Laminin.

**Figure 2.**
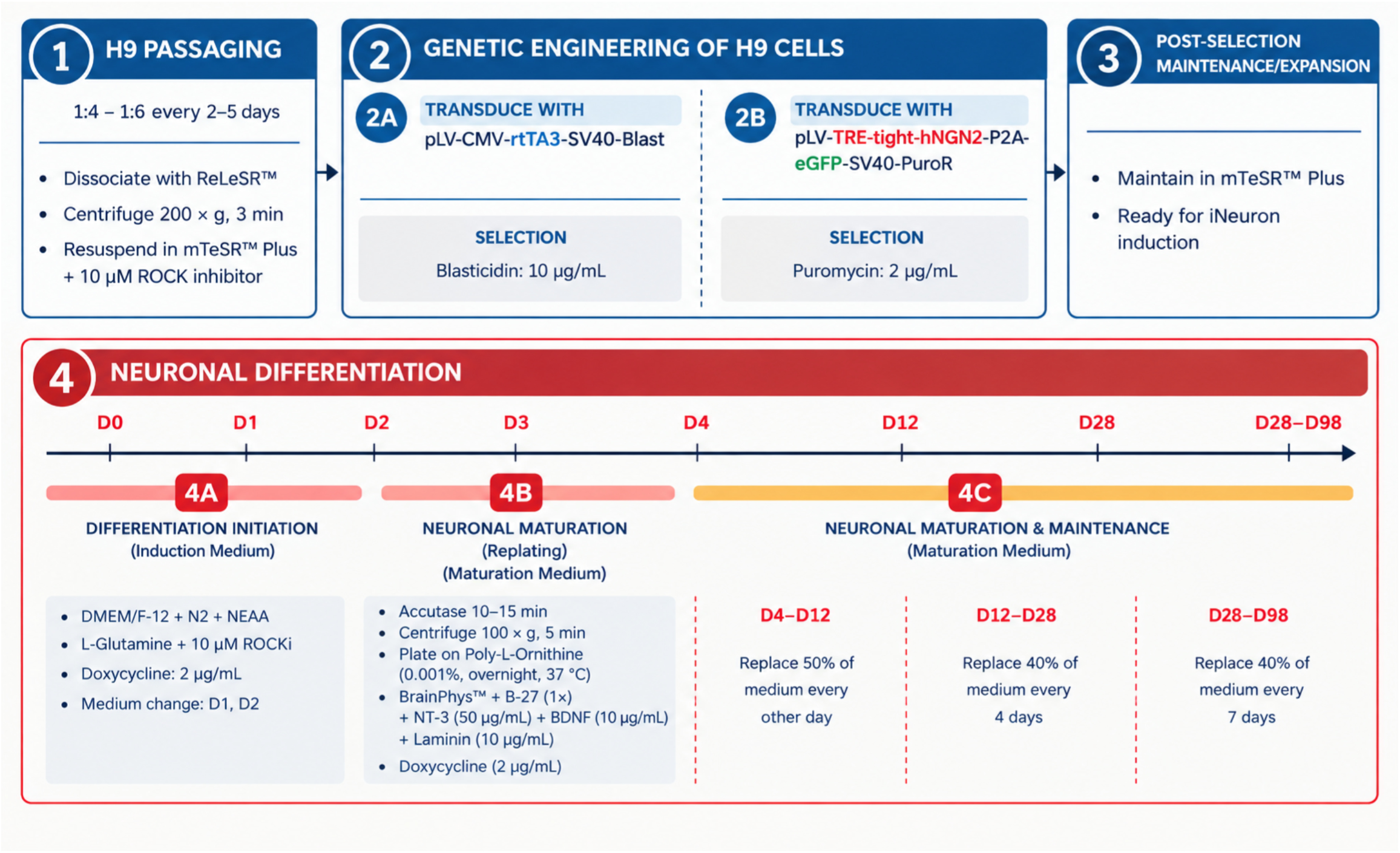
Summary of neuronal differentiation with the pLV-TetO-hNGN2-P2A-eGFP-SV40-PuroR strategy. H9 cells were stably maintained (Step 1), followed by transduction with rtTA3 and pLV-TetO-hNGN2-P2A-eGFP-SV40-PuroR (Step 2). Cells were allowed to recover from transduction and maintained until they resumed normal proliferation and passaging frequency (Step 3). Puromycin-selected cells were treated with induction medium to initiate differentiation (Step 4A) for 3 days; on day 3, cells were replated in maturation medium (Step 4B). During the subsequent maintenance stage, neurons were kept with medium changes at defined intervals (Step 4C).

Using this protocol, we observed individual cells extending neurites by 3 days post-differentiation (DPD3)(Fig 3A). After replating differentiation-initiated cells into maturation medium, more pronounced neurite outgrowth was observed while cells with extended processes remained largely non-clustered (DPD4 to DPD32). Between DPD32 and DPD60, cells began forming soma–soma clusters with radially extending neurites, a morphology consistent with progressive neuronal maturation (Hörberg et al., 2022).

**Figure 3.**
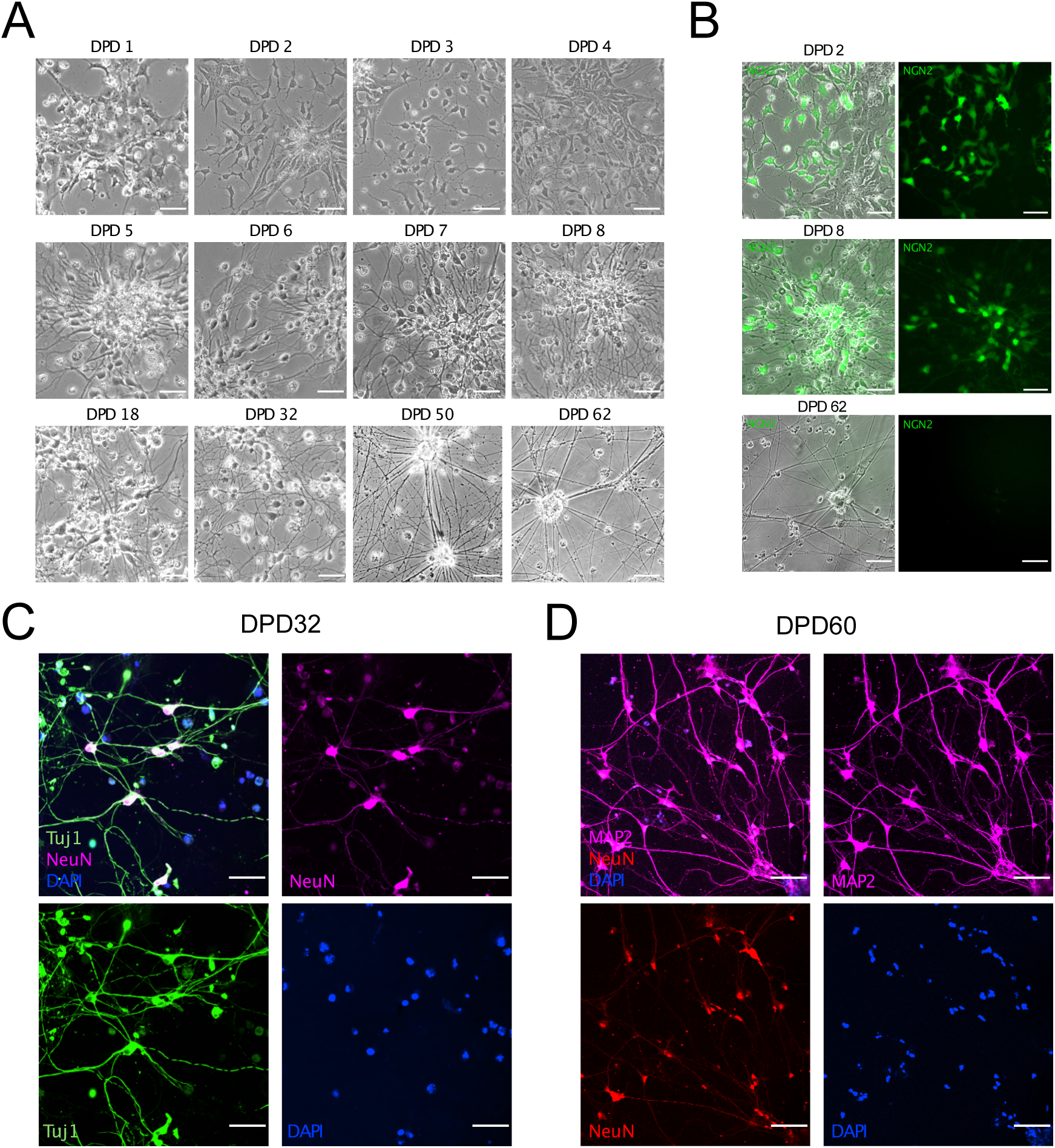
Validation of neuronal morphology and functional markers during differentiation. (3A) Light-phase microscopy images of hNGN2-induced iNeurons over time starting 1 day post-differentiation (DPD) to DPD62. (3B) hNGN2-GFP signal at various time points post-differentiation. Green-channel images indicate reconstituted hNGN2-GFP expression. (3C, D) DPD32 and DPD60 differentiated neurons immunostained for the neuronal marker Tuj1 and the mature neuronal markers NeuN and MAP2. Scale bar = 20 μm.

With the pLVTetO-hNGN2-P2A-eGFP SV40-PuroR construct, GFP is expressed through an IRES element downstream of NGN2. After doxycycline addition, we observed GFP-positive cells beginning at DPD2, indicating expression of the transgene (Fig. 3B). GFP signal remained readily detectable through DPD8 but was markedly diminished by DPD62. This progressive loss of transgene expression may reflect differentiation-associated transgene silencing during neuronal maturation, consistent with recent evidence showing that conventional transgenes are frequently silenced during pluripotent stem cell differentiation, including neuronal differentiation (Uenaka et al., 2026). Expression of the immature neuronal marker Tuj1 and the mature marker NeuN was detected by DPD30, and the functional/mature markers MAP2 and NeuN were present by DPD60 (Fig. 3C, D).

Using this optimized protocol, we achieved >50% neuronal yield with no evidence of residual proliferating cells. We did observe a population of non-dividing, neurite-negative cells with reduced cell size and positive propidium iodide (PI) staining without permeabilization (Supplementary Fig. 1, 29.1 ± 6.6% of neuronal population). We hypothesize that this cell debris arises from heterogeneous NGN2 expression levels or insufficient differentiation signaling strength.

### Gene expression analysis confirms neuronal identity

To confirm neuronal identity and culture homogeneity, cells were collected at multiple differentiation stages for RT-qPCR and RNA-seq analysis (Fig. 4). The stemness marker SOX2 and neural stem cell marker Nestin were progressively downregulated through DPD35 compared to the starting H9 cell stem cell population (Fig. 4A) while the oligodendrocyte marker OLIG2 was expressed at early stages but declined by DPD35. PAX6, a marker of dorsal telencephalic progenitor identity (Quinn et al., 2007), was strongly upregulated by DPD20, coinciding with a marked increase in MAP2 expression, consistent with progression toward a mature neuronal phenotype.

**Figure 4.**
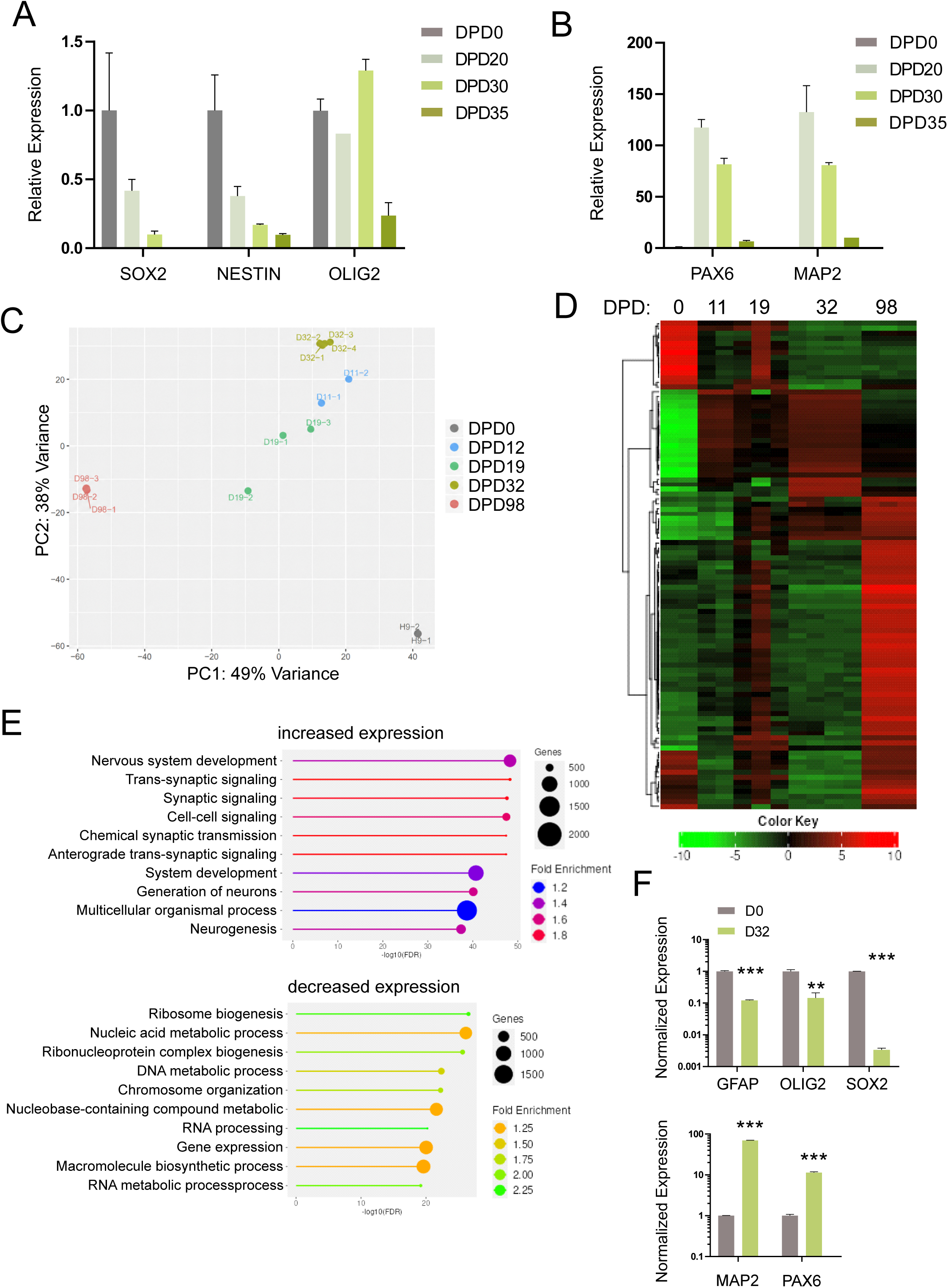
Gene expression analysis of differentiation time course. (4A, B) qPCR of stemness and neuronal markers at different stages of differentiation. Expression fold-change of SOX2, NESTIN, and OLIG2 in undifferentiated H9 cells (DPD0) versus DPD20, DPD30, and DPD35 iNeurons; PAX6 and MAP2 indicate neuronal maturation in the respective groups. Expression was normalized to the TXN2 gene, N=3. Missing values were excluded from the analysis. (4C) PCA plot of RNA-seq data showing transcriptomic variation among cells from DPD0 to DPD98. Each point represents a sample, plotted along PC1 and PC2, which together capture the largest sources of variance in gene expression. Samples cluster according to differentiation stage. (4D) Heatmap of transcriptomic differences across groups after unsupervised hierarchical clustering. The top 200 differentially expressed genes are shown. Image generated with iDEP 2.50. (4E) Up- and downregulated pathways in DPD32 vs. DPD0. The size of each lollipop corresponds to the number of genes in that pathway; color indicates the fold-enrichment level. Enrichment analysis performed using GO Biological Process terms. Image generated with iDEP 2.50. (4F) Normalized RNA-seq expression from the samples in Fig. 4C. Shown are the glial marker GFAP, the oligodendrocyte marker OLIG2, the stem cell marker SOX2, and the neuronal markers MAP2 and PAX6. N=2 for D0 and N=4 for D32. Statistical analysis was performed using an unpaired t-test. **p<0.01, ***p<0.005.

Principal component analysis (PCA) of RNA-seq data showed clear separation between undifferentiated (DPD0), intermediate (DPD11-DPD32), and late-stage (DPD98) samples (Fig. 4C), and hierarchical clustering revealed distinct transcriptomic profiles across these time points (Fig. 4D). Differential expression analysis between DPD0 and DPD32 identified a robust set of the top 300 differentially expressed genes (DEGs) driving this separation (Fig. 4E). KEGG pathway analysis of these DEGs showed that more than half of the top 10 enriched pathways were associated with neurogenesis, while the most significantly downregulated pathways were linked to DNA metabolism and ribosome biogenesis — consistent with a transition from a proliferative stem cell state to a post-mitotic neuronal state. RNA-seq analysis of non-differentiated H9 cells and DPD32 iNeurons revealed marked decreases in progenitor/glial-associated markers, including GFAP (∼8-fold), OLIG2 (∼7-fold), and SOX2 (∼300-fold). In contrast, neuronal markers MAP2 and PAX6 were increased in iNeurons, with MAP2 showing a dramatic 70-fold increase (Fig. 4F).

### Calcium-induced membrane depolarization assay

To implement a functional assay for neuronal activity, we combined optogenetic stimulation with calcium imaging in H9-derived neurons cultured on coverslips (Fig. 5A). Neurons were co-transduced with hSyn-ReaChR-citrine (Lin et al., 2013a)., encoding the red-shifted channelrhodopsin ReaChR fused to citrine in differentiated neurons, and MAP2-GCaMP3, a genetically encoded calcium indicator driven by the neuron-specific MAP2 promoter. Citrine fluorescence identifies transduced neurons, while GCaMP3 fluorescence reports intracellular calcium flux associated with membrane depolarization. ReaChR was activated with 630 nm light, and GCaMP3 was excited at 480–500 nm with emission collected at 510–520 nm, allowing spectral separation of stimulation and readout channels. All imaging was performed at 37 °C to preserve neuronal viability.

**Figure 5.**
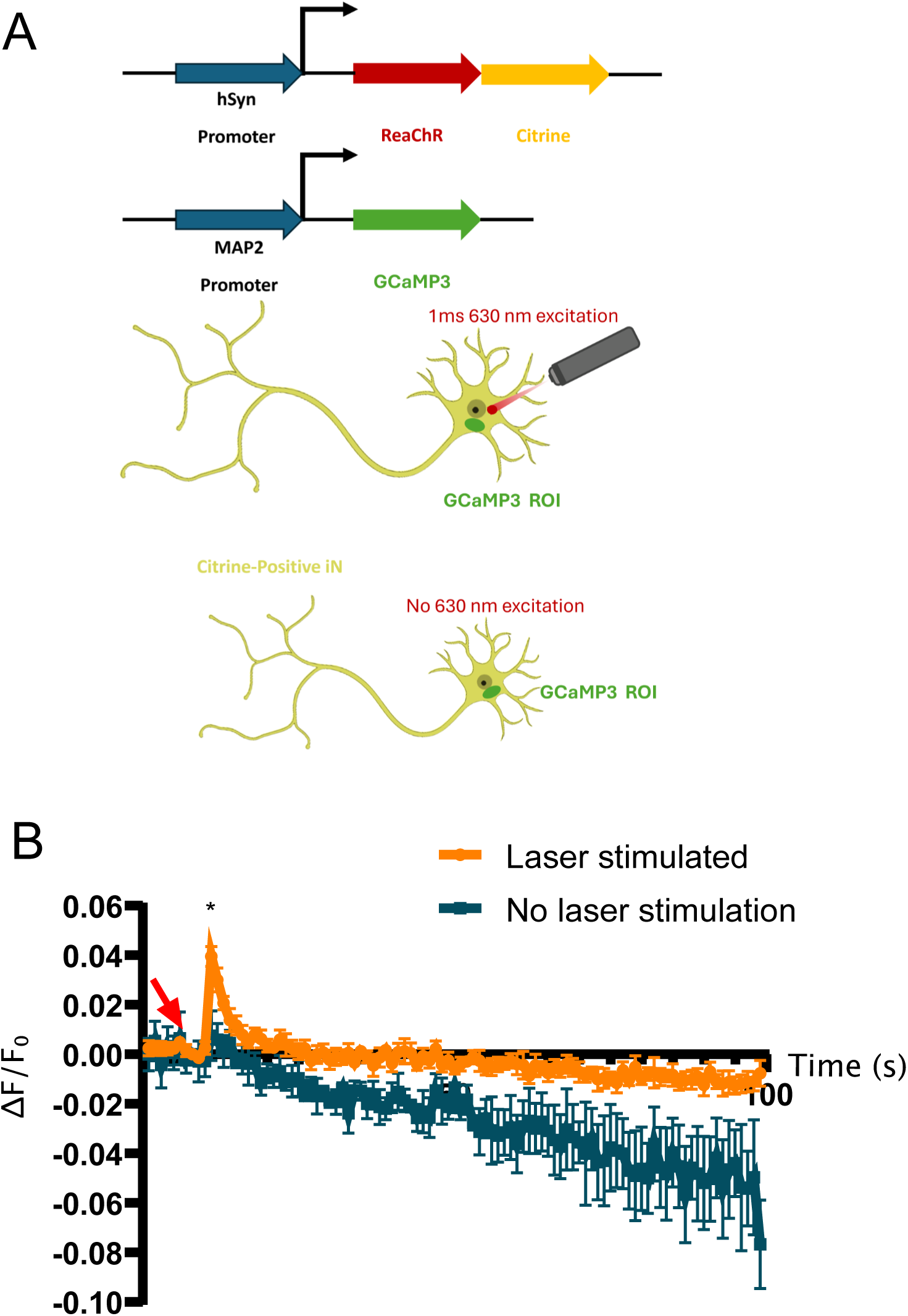
Live-cell assay for optogenetic neural excitation in induced neurons. (5A) Schematic of the optogenetic stimulation and calcium-imaging setup. In transduced iNeurons, expression of the red-shifted variant of channelrhodopsin ReaChR as well as the calcium indicator GCaMP3 expression are driven by the hSyn and MAP2 human neuronal promoters, respectively. Laser-stimulated neurons were live-imaged for GCaMP3 fluorescence. Only citrine-positive neurons were scored. (5B) GCaMP3 ΔF/F fluorescence traces of stimulated and non-stimulated iNeurons. Citrine-positive cells were laser-stimulated for 1 ms. GCaMP signal was scored only in the cell body throughout the experiment, excluding the laser-stimulated area. Red arrow indicates the laser stimulation. Asterisks indicate statistical significance p < 0.05 by paired t-test at the time of stimulation (N=4–5).

Brief (1 ms) laser stimulation of individual cell bodies elicited a corresponding increase in GCaMP3 signal, while neighboring, non-stimulated neurons showed no change in fluorescence (Fig 5B). These results demonstrate a simple, light-based method for assessing the functionality of differentiated neurons *in vitro*.

Here we established an optimized NGN2-induction protocol that generates pure, glial-free human neuronal cultures from H9 stem cells at high yield (>50%), overcoming the low viability by uncoupling antibiotic selection from NGN2 induction and avoiding the use of DNA damaging nucleoside analogs. The resulting neurons mature normally, both morphologically and molecularly, and are functionally active. We confirmed this using a simple optogenetic and calcium imaging assay that provides fast, single-cell readouts of neuronal activity. Together, this scalable differentiation protocol and simple functional assay offer an easy, low-barrier platform for producing and validating homogeneous, functional human neurons for molecular and omics studies.

## Materials and Methods

### Gene expression constructs

The doxycycline-inducible pLV-TetO-hNGN2-eGFP-Puro plasmid was a gift from Kristen Brennand (Addgene plasmid #79823). A derivative lacking the PuroR cassette, pTP5693 (pLV-TetO-hNGN2-P2A-eGFP-STOP), was generated by removing the T2A-PuroR sequence from the pLV-TetO-hNGN2-eGFP-Puro and introducing a stop codon. The final pLV-TetO-hNGN2-P2A-eGFP-SV40-PuroR construct (pTP5716) was subsequently generated by Gibson assembly of the pTP5693 backbone with an SV40-PuroR fragment amplified from pBABEpuro GFP-LC3 (Addgene plasmid #22405)(Fung et al., 2008), a gift from Jayanta Debnath. The pLenti CMV rtTA3 Blast (w756-1) plasmid was a gift from Eric Campeau (Addgene plasmid #26429) and was used as the source of the CMV-driven rtTA3 expression cassette. pLenti-ReaChR-Citrine was a gift from Roger Tsien (Addgene #50956)(Lin et al., 2013b), and MAP2-GCaMP3 was a gift from John Gearhart (Addgene #43917)(Addis et al., 2011). NGN2-AID, ERT2, and FKBP constructs were made using the lenticrispr v2 backbone (Addgene plasmid #52961) after deletion of Cas9; details available upon request.

### H9-iNeuron Differentiation

H9 human embryonic stem cells (hESCs; gift from Jonghwan Kim) were maintained on Matrigel-coated culture vessels in mTeSR™ Plus medium (STEMCELL Technologies) with daily medium replacement. Culture surfaces were coated with Corning® Matrigel® diluted 1:100 in DMEM/F-12 media (Gibco, Thermo Fisher Scientific) and incubated at 37°C for 30 min before use.

H9 cells were passaged every 2–5 days at a 1:4–1:6 ratio. Cells were dissociated using ReLeSR™ and collected by centrifugation at 200 × g for 3 min. Cells were resuspended in mTeSR™ Plus supplemented with 10 µM Y-27632 dihydrochloride (MedChemExpress) and replated onto vessels coated with Matrigel® hESC-Qualified Matrix, LDEV-Free (Corning). For cryopreservation, cells were resuspended in FreSR™ Cryopreservation Medium (STEMCELL Technologies) and stored initially at −80°C before transfer to liquid nitrogen or −150°C for long-term storage.

To prepare for neuronal induction, H9 cells were transduced with lentiviral vectors at 20%–30% confluency. Concentrated viral preparations were generated by ultracentrifugation at 48,000 × g for 1.5 h, sterilized with VWR 0.45 µm PES Syringe Filters (Avantor) and resuspended in mTeSR™ Plus. Cells were treated with Polybrene (6 µg/mL) for 15 min before viral exposure. One-quarter of the viral preparation was added on day 1 and the remaining three-quarters on day 2. Cells were subsequently maintained or passaged according to confluency. pLenti-CMV-rtTA3-Blast was transduced first, selected with blasticidin (10 µg/mL) then transduced with pTP5716, and selected with puromycin (1–2 µg/mL). Cells were expanded and stocked at this stage for later use.

### iNeuron Differentiation and Culture

H9 cells were dissociated and plated at a 1:2.5 to 1:3 ratio on Matrigel-coated plates in induction medium composed of DMEM/F-12 (Gibco™) supplemented with 1× N2 supplement ((Gibco™)), 1× Non-Essential Amino Acid (NEAA) Solution (Cytiva), 1× L-Glutamine (Gibco™), 10 □µM Y-27632 dihydrochloride, and 2 □µg/mL doxycycline (Fisher Scientific). On the second day, cultures were gently washed once with 1× PBS and replenished with fresh induction medium. Medium was changed again on Day 3. On Day 4, cells were passaged for neuronal maturation.

For maturation, plates were pre-coated with 0.001% Poly-L-Ornithine in PBS (Sigma-Aldrich) and incubated overnight at 37□ °C. On Day 4, induced H9 cells were washed once with 1× PBS, treated with pre-warmed Accutase for 10–15 minutes, and resuspended in 3× mTeSR™ Plus medium. After centrifugation at 100□ ×g for 5 minutes, cells were resuspended in maturation medium consisting of BrainPhys™ Neuronal Medium (STEMCELL Technologies) supplemented with 1× B-27™ Supplement (Thermo Fisher Scientific), 10 ng/mL Recombinant human NT-3 (PeproTech), 10 □ng/mL Recombinant Human BDNF Protein, CF (R&D System), 1 µg/mL Laminin (MilliporeSigma), and 2□ µg/mL doxycycline. Cells were incubated at 37□ °C. To maintain neuronal viability, exposure to ambient conditions was limited to under 5 minutes during medium changes.

Maturation medium was refreshed by replacing 50% of the volume every other day for the first 4 days. After Day 12, medium was changed every 4 days by removing 40% of the existing medium and adding 60% fresh medium. After Day 28 of differentiation, medium changes were performed every 7 days. Gentle pipetting and minimal mechanical disturbance were essential to avoid neuronal detachment and death.

### qRT-PCR analysis

Total RNA was isolated from cultured cells by scraping the cells directly into PBS without trypsinization, followed by RNA purification using the Monarch RNA Cleanup Kit (NEB) according to the manufacturer’s instructions. cDNA was synthesized from the purified RNA using the SuperScript IV First-Strand Synthesis System (Thermo Fisher Scientific) according to the manufacturer’s instructions. qRT-PCR was performed using PowerUp SYBR Green Master Mix (Thermo Fisher Scientific) on a LightCycler 480 Real-Time PCR System (Roche).

The primer sequences used for amplification of SOX2, PAX6, NESTIN, MAP2, GFAP, OLIG2, and TXN2 were obtained from Liu et al. (2018)(Liu et al., 2018)and were as follows: SOX2, Fwd 5′-TACAGCATGTCCTACTCGCAG-3′, Rev 5′-GAGGAAGAGGTAACCACAGGG-3′; PAX6, Fwd 5′-ATGTGTGAGTAAAATTCTGGGCA-3′, Rev 5′-GCTTACAACTTCTGGAGTCGCTA-3′; NESTIN, Fwd 5′-GGCGCACCTCAAGATGTCC-3′, Rev 5′-CTTGGGGTCCTGAAAGCTG-3′; MAP2, Fwd 5′-GATGGCTTCAGGGCTAAACA-3′, Rev 5′-GCTTTCAAGGGACTCCATGGT-3′; GFAP, Fwd 5′-TCATCGCTCAGGAGGTCCTT-3′, Rev 5′-CTGTTGCCAGAGATGGAGGTT-3′; and OLIG2, Fwd 5′-GGACAAGCTAGGAGGCAGTG-3′, Rev 5′-ATGGCGATGTTGAGGTCGTG-3′.

Gene expression was normalized to TXN2, using the following primers: Fwd 5′-ACCACACAGACCTCGCCATT-3′ and Rev 5′-AGGCCTCCAACTGATCCTCAT-3′. Relative gene expression was calculated using the ΔΔCt method (Schmittgen and Livak, 2008).

### RNAseq and differential gene expression analysis

250 ng RNA from differentiated neurons was used to prepare mRNA with the NEBNext Poly(A) mRNA magnetic isolation kit (NEB E7490S). 192 ng mRNA was used to prepare sequencing libraries with the NEBNext UltraExpress RNA Library Prep Kit (NEB E3330S) which were sequenced using dual index primers via the NovSeq X plus platform (PE150)(Novogene). Analysis of raw data was performed using FastP for pre-processing (Chen et al., 2018), HISAT2 alignment to the hg38 build of the human genome (Kim et al., 2019), and DESeq2 for quantitation of differentially expressed genes (Love et al., 2014). The PCA plot was generated using R in Galaxy EU. Heatmap analysis was performed using hierarchical clustering in iDEP 2.50 (iDEP). Enrichment analysis was performed using Gene Ontology (GO) Biological Process terms in iDEP 2.50.

### Quantification of PI-positive non-neuronal cells

To quantify non-dividing, neurite-negative cells and cellular debris, 1× propidium iodide (PI) was added directly to live cell cultures without fixation or permeabilization. Cells were imaged using a Nikon ECLIPSE Ts2-W inverted microscope with a 20× objective under TRITC/Cy3 fluorescence illumination (560-nm excitation). Six representative fields were acquired for each condition. PI-positive cells were manually counted in each field and expressed as a percentage of the total cell population. Cells exhibiting reduced cell size, absence of neurite outgrowth, and positive PI fluorescence were classified as the non-dividing, neurite-negative population.

### Immunostaining

Immunostaining was performed based on a previously published protocol for neuronal cultures (Yin et al., 2023). H9-derived neurons were briefly washed with 1× PBS and fixed with 4% paraformaldehyde in 1× PBS for 10 min at room temperature. Cells were permeabilized with 0.7% Triton X-100 in 1× PBS for 15 min at room temperature, followed by three washes with PBS. Cells were incubated overnight at 4°C with primary antibodies diluted in antibody buffer containing 1% BSA, 5% horse serum (Thermo Fisher Scientific), and 0.1% Triton X-100 in 1× PBST. Primary antibodies included mouse anti-MAP2 (1:600–1:1000; Novus Biologicals, NBP1-92711SS), mouse anti-Tuj1 (1:400; Fisher Scientific, MAB1195-SP), and rabbit anti-NeuN (1:400; GeneTex, GTX132974). Following three PBS washes, cells were incubated with goat anti-mouse Alexa Fluor 647 (Thermo Fisher Scientific) and donkey anti-rabbit Alexa Fluor 594 (Thermo Fisher Scientific) secondary antibodies (1:400) together with Hoechst 33342 (1:3000, Fisher Scientific) for 1 h at room temperature in 7% BSA/PBS. Cells were then washed three times with PBS and mounted using ProLong Gold Antifade Mountant (Thermo Fisher Scientific, P10144). Coverslips were allowed to dry overnight at room temperature protected from light.

### Optogenetic stimulation and calcium imaging

H9-derived neurons cultured on coverslips were co-transduced with hSyn-ReaChR-Citrine and MAP2-GCaMP3 lentivirus as described above. MAP2-GCaMP3 expresses the genetically encoded calcium indicator GCaMP3 under the control of the human MAP2 promoter. ReaChR-Citrine was used for optogenetic stimulation, and GCaMP3 fluorescence was used to monitor neuronal calcium responses. Live-cell imaging was performed at 37°C using a Nikon AX R confocal microscope equipped with an NSPARC detector. Citrine-positive neurons were identified before stimulation and were included for analysis. ReaChR was activated using far-red 630-nm laser illumination, and GCaMP3 fluorescence was detected using green 488-nm excitation with emission collected at 510–520 nm. Individual neuronal cell bodies were stimulated with 1-ms laser pulses, and changes in GCaMP3 fluorescence were recorded before and after stimulation. Changes in fluorescence were quantified as ΔF/F using ImageJ, with F representing the baseline fluorescence before stimulation. Statistical significance p < 0.05 was determined by paired t-test (N=4).

## Key resources table

| **Recombinant DNA** | **Source** | **Identifier** |
|---|---|---|
| AID-NGN2 in lentiCRISPR v2 (Puro), GFP-free | This work | pTP5402 |
| EF1α-AID-linker-hNGN2-P2A-Puro in lentiCRISPR v2 | This work | pTP5592 |
| EF1α-ERT2-linker-hNGN2-P2A-Puro in lentiCRISPR v2 | This work | pTP5593 |
| EF1α-hNGN2-linker-ERT2-P2A-Puro in lentiCRISPR v2 | This work | pTP5594 |
| EF1α-NGN2-linker-NLS-FKBP-V in lentiCRISPR v2 | This work | pTP5690 |
| pLV-TetO-hNGN2-P2A-eGFP-STOP | This work | pTP5693 |
| pLV-TetO-hNGN2-P2A-eGFP-SV40-PuroR | This work | pTP5716 |
| pBABEpuro GFP-LC3 | Fung et al., 2008(Fung et al., 2008) | Addgene #22405 |
| pLV-TetO-hNGN2-eGFP-Puro | Ho et al., 2015(Ho et al., 2016) | Addgene #79823 |
| pLenti CMV rtTA3 Blast (w756-1) | Unpublished | Addgene #26429 |
| pLenti-ReaChR-Citrine | Lin et al., 2013(Lin et al., 2013c) | Addgene #50956 |
| MAP2-GCaMP3 | Addis et al., 2011(Addis et al., | Addgene |
|  | 2011) | #43917 |

## Acknowledgements

We are grateful to members of the Paull laboratory for their assistance and comments on the manuscript. The work was supported by NIH R01 NS126747 to T.P.

