## Supplemental Figure 1 for "Efficient NGN2-induced neuronal differentiation from human stem cells for DNA repair studies and functional analysis"

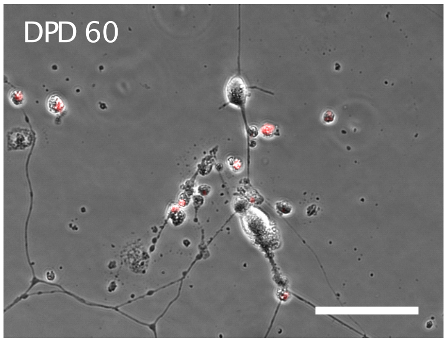


**Supplementary Fig. 1 — Presence of PI-positive cell debris**
Live neurons with cell debris are shown, stained with propidium iodide (PI) without permeabilization.
Based on N=6 images scored, 29.1 ± 6.6% of cells in the culture were PI-positive (dead). Scale bar = 20 μm.
